# Two homologous Alt a1-like fungal proteins possess dual activities in HIR-associated immune signaling and EDS1-dependent cell death

**DOI:** 10.64898/2026.08.11.744170

**Authors:** Tobias Müller, Marat Magomedov, Charlene Chaudy, Matthias Hahn, David Scheuring

## Abstract

Necrotrophic fungi secrete numerous Cell Death-Inducing Proteins (CDIPs) that manipulate host immunity to promote disease, yet the signaling pathways underlying their phytotoxic activity remain poorly understood. Here, we identify the *Botrytis cinerea* Hypersensitive response-inducing protein 1 (Hip1) as a close homolog of the recently described *Sclerotinia sclerotiorum* effector Plant Early Immunosuppressive Effector 1 (PEIE1) and investigate the molecular basis of its activity. HIP1 and PEIE1 share high sequence similarity and a conserved AlphaFold-predicted Alt a1-like fold, they interact with the Arabidopsis plasma membrane protein HIR4, and they induce strong necrosis in *Nicotiana benthamiana*. Despite their high structural similarity, Hip1 and PEIE1 differ in their reported roles during fungal infection. Unexpectedly, Hip1-induced cell death requires the central immune regulator ENHANCED DISEASE SUSCEPTIBILITY 1 (EDS1) as well as the downstream helper NLR network comprising ADR1 and NRG1. Together, our findings establish Hip1 as a closely related homolog of PEIE1 and suggest that these closely related Alt a1-like proteins possess dual activities: modulation of HIR-associated immune signaling and activation of EDS1-dependent host cell death.

## Introduction

Plant pathogens cause significant annual crop losses. The socioeconomically important fungal pathogen *Botrytis cinerea* (*Botrytis* hereafter), causing grey mold rot, has a broad host range and infects over 1000 plant species, including economically important vegetables (e.g. tomato, lettuce and cucumber), fruit crops (e.g. grapevine and strawberry) and ornamental flowers (e.g. rose and gerbera) (Fillinger and Elad, 2016). The life cycle of this necrotrophic fungus involves rapid killing of plant cells, colonization of dead host tissue and ultimately massive reproduction via conidiospores (Bi et al., 2022). To overcome plant barriers initially, *Botrytis* employs a combination of high mechanical force (Müller et al., 2024) and secretion of toxic proteins and metabolites, effectors and cell-wall degrading enzymes (Espino et al., 2010; González-Fernández et al., 2015; Bi et al., 2022).

Although substantial progress has been made in understanding the infection process of *Botrytis* in the last decade (Bi et al., 2022), the function of numerous secreted proteins is still unclear and more and more different modes of plant cell death are discovered: besides cell wall-degrading enzymes (CWDEs), such as endopolygalacturonases (Have et al., 1998; Kars et al., 2005), and cytolytic proteins of the NEP1-like protein (NLP) class (Seidl and van den Ackerveken, 2019; Pirc et al., 2023), more recent evidence points to sophisticated means targeting plant immune signaling. Among fungal proteins associated with plant immunity, the Alt a1 protein family has attracted increasing attention. The *Alternaria alternata* protein Alt a1 can induce plant defence responses and cell death when transiently expressed in *N. benthamiana* (Zhang et al., 2025). The phytotoxicity of most so-called Cell Death Inducing Proteins (CDIPs) seems to rely on receptor recognition at the cell surface as pathogen-associated molecular patterns (PAMPs). In case of *Botrytis* CDIPs, this induced plant PAMP-triggered immunity (PTI) often involves the strongest form of plant defence, the hypersensitive response (HR) which includes programmed cell death of infected tissue (Boutrot and Zipfel, 2017). Since HR is usually associated with plant effector-triggered Immunity (ETI), plant cell death triggered by activation of PTI seems to be a specific feature of necrotrophic pathogens. Notably, it has been reported for the first time recently that *Botrytis* secretes also a cell death inducing protein which functions inside plant cells. Congo red hypersensitivity 1 (Crh1) is secreted into the plant apoplast and later translocated into plant cells where it induces plant cell death (Liang et al., 2026; Bi et al., 2021).

In many cases, recognition of fungal cell death-inducing proteins (CDIPs) at the plant cell surface depends on the co-receptors BRI1-ASSOCIATED KINASE 1 (BAK1) and SUPPRESSOR OF BIR1-1 (SOBIR1), which associate with ligand-binding receptor-like proteins (RLPs) to initiate immune signaling) (Schellenberger et al., 2019). A well-characterized example is the perception of Botrytis endopolygalacturonases by the RLP RESPONSIVENESS TO BOTRYTIS POLYGALACTURONASES 1 (RBPG1), which requires both SOBIR1 and BAK1 for downstream signaling (Zhang et al., 2014; Zhang et al., 2021). Similarly, the *Botrytis* CDIPs Xyloglucanase 1 (XYG1Cell Death-Inducing protein 1 (CDI1), and the glycoside hydrolase Gas5A depend on BAK1 and SOBIR1 for full activity, although their cognate receptors remain enigmatic (Zhu et al., 2017; Zhu et al., 2023; Müller et al., 2026). In contrast, the Botrytis CDIP Hypersensitive Response-Inducing Protein 1 (Hip1) induces cell death independently of BAK1 and SOBIR1, indicating that it engages a mechanistically distinct signaling pathway (Jeblick et al., 2023). While the plant receptor for Hip1 has remained elusive, its orthologue Plant Early Immunosuppressive Effector 1 (PEIE1) from *Sclerotinia sclerotiorum* (*Sclerotinia* hereafter) was recently shown to interact with members of the Hypersensitive Induced Reaction (HIR) protein family, identifying the first candidate host target for this class of fungal proteins (Liu et al., 2024). In general, accumulation of HIR proteins upon pathogen attack results in activation of host HR (Zhou et al., 2010; Jung and Hwang, 2007). By inhibiting HIR oligomerization, PEIE1 dampens plant immunity and serves as virulence factor for *Sclerotinia* (Liu et al., 2024). Although many individual *Botrytis* CDIP mutants display little or no virulence phenotype (Leisen et al., 2022), our data reveal that Hip1 makes a measurable contribution to virulence during Arabidopsis infection. Notably, the potential phytotoxicity of PEIE1, which is a key feature of its orthologue Hip1, was not addressed in the work from Liu et al. (Liu et al., 2024).

In the present study, we report on the function of *Botrytis* Hip1 and its *Sclerotinia* orthologue PEIE1. We show that Hip1 functions as virulence factor for *Botrytis* infection of Arabidopsis. Furthermore, we demonstrate that Hip1 from *Botrytis* and PEIE1 from *Sclerotinia* are genetically, structurally and functionally highly similar and show that their phytotoxic activity depends on intracellular plant immune signaling. We identify the plant immune regulator ENHANCED DISEASE SUSCEPTIBILITY 1 (EDS1) as a crucial component required for Hip1-and PEIE1-induced cell death, revealing that extracellular fungal cell death-inducing proteins can engage EDS1-dependent immune signaling through a pathway that is genetically distinct from the canonical SOBIR1/BAK1-dependent LRR-RP pathway (Fliegmann et al., 2026).

## Results

### Hip1 and PEIE1 share sequence similarity and a conserved Alt a1-like fold

Recently, Plant Early Immunosuppressive Effector 1 (PEIE1) was identified as a secreted effector of *Sclerotinia sclerotiorum* (Liu et al., 2024). Its homolog in *Botrytis cinerea*, Hypersensitive response-inducing protein 1 (Hip1), was previously characterized as a cell death-inducing protein; however, the underlying mechanism remains unclear (Jeblick et al., 2023). AlphaFold-based structural predictions and sequence comparisons revealed a high degree of similarity between Hip1 and PEIE1 (Fig. 1), suggesting that these proteins may share related functions. Interestingly, although the *Botrytis* protein Hip2 adopts a similar overall fold, it is more divergent from Hip1 than PEIE1 is (Fig. 1B), highlighting the close relationship between Hip1 and PEIE1.

**Fig. 1:**
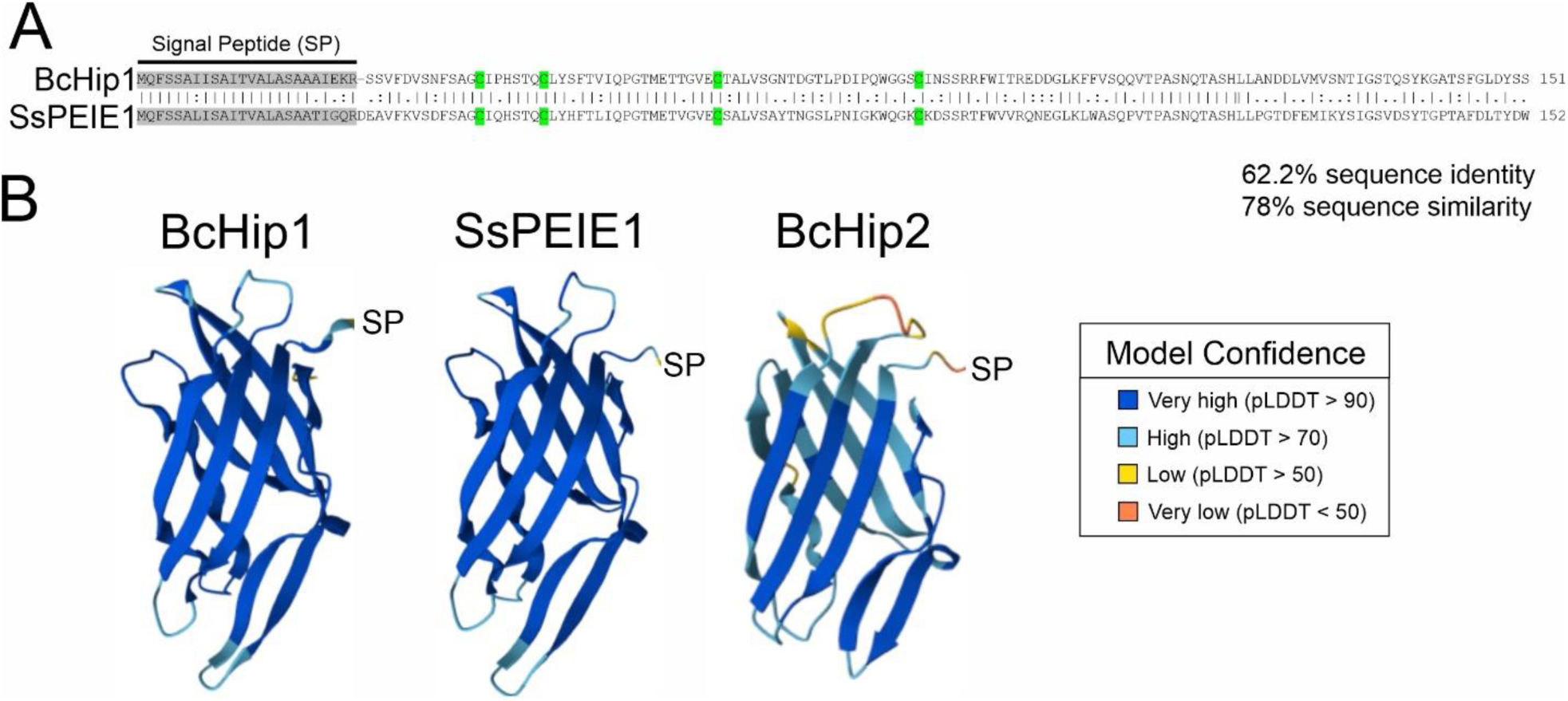
BcHip1 and SsPEIE1 are orthologs. A) Sequence alignment of BcHip1 and SsPEIE1. Identical amino acids are indicated with a vertical line, similar amino acids with two dots and non-related amino acids with one dot. Grey background symbolizes predicted signal peptides (SignalP 5.0). Conserved cysteine residues are labeled in green. B) AlphaFold predictions of BcHip1, SsPEIE1 and BcHip2. Confidence level is shown as predicted by Local Distance Difference Test (LDDT) and color-coded as indicated.

### Hip1 and PEIE1 interact with the plant PM protein HIR4

To investigate whether Hip1 functions similarly to PEIE1 by targeting plasma membrane-localized Hypersensitive Induced Reaction (HIR), as recently described for PEIE1 (Liu et al., 2024), we performed yeast interaction assays using the mating-based split ubiquitin system (mbSUS) (Grefen et al., 2009). Among the four Arabidopsis HIR proteins (HIR1–HIR4), PEIE1 was previously shown to interact with HIR2 and, more strongly, HIR4, thereby suppressing plant immunity (Liu et al., 2024). Consistent with this finding, mbSUS assays revealed a specific interaction between Hip1 and Arabidopsis HIR4, whereas the closely related protein Hip2 showed no detectable interaction (Figure 2). PEIE1, included as a positive control, also interacted with HIR4.

**Fig. 2:**
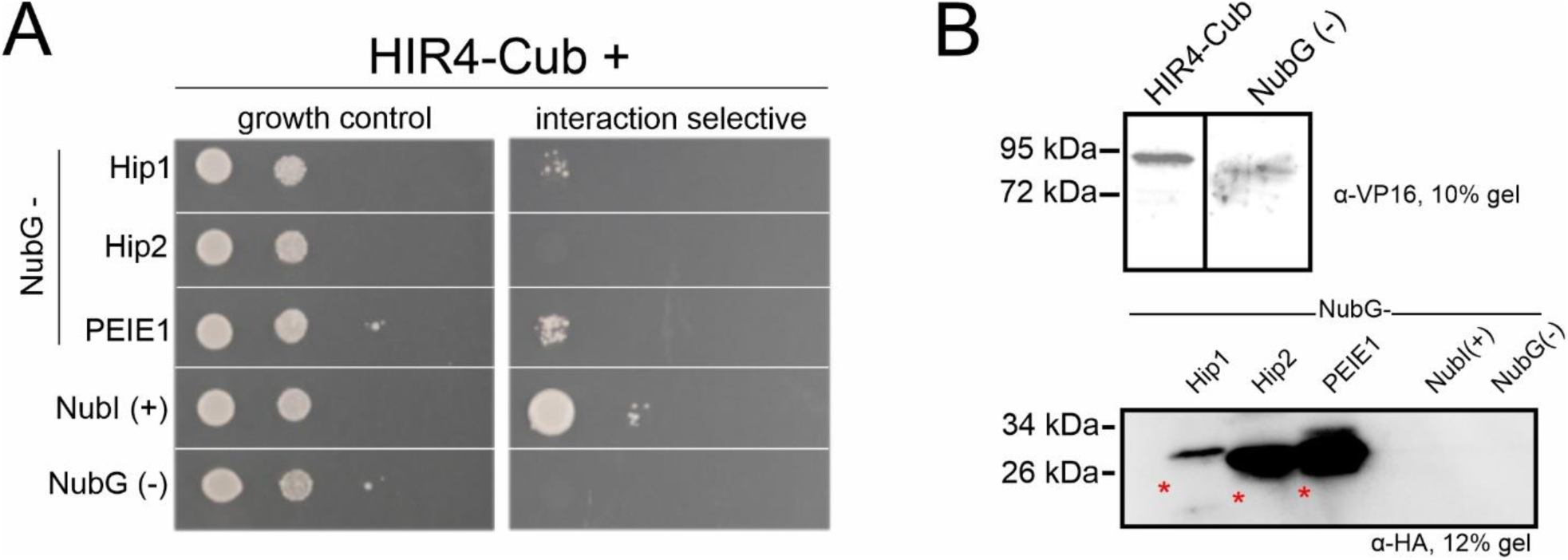
Hip1 and PEIE1 interact with Arabidopsis HIR4. A) Yeast mating-based Split-Ubiquitin System (mbSUS) growth assays for testing interaction of HIR4-Cub with NubG-Hip1, NubG-Hip2 and NubG-PEIE1. Yeast strain THY.AP4 (MATa) was transformed with HIR4-Cub and mated with THY.AP5 (MATα) strains that were transformed with NubG-fusions. Growth of mated yeasts was assessed in dilution series of OD_600_ from 1 to 0.001 at 30°C. Growth control media (CSM +adenine/histidine) was used to confirm yeasts maintaining fusion constructs. Interaction selective media (CSM + 5 µM methionine) was used for analyzing interaction-dependent activation of reporter genes. NubI wild type was used as positive control, NubG as negative control. After 48h, interaction-dependent growth was detected for Hip1 and PEIE1 with HIR4. B) Expression controls via Western Blot show expression of HIR4-Cub and all NubG-fusion constructs. Expected sizes are indicated by an asterisk. The experiment was performed three times. One representative experiment is shown.

Taken together, these results demonstrate that Hip1 and PEIE1 share structural similarity and host interaction targets.

### Hip1 contributes to *Botrytis* virulence on Arabidopsis in a HIR4-dependent manner

Since PEIE1 has been identified as a secreted virulence factor in *Sclerotinia* (Liu et al., 2024), we investigated whether its *Botrytis* homolog Hip1 similarly contributes to fungal pathogenicity. To this end, we compared disease development caused by *Botrytis* wild type (B05.10) and a previously established *hip1* knockout mutant (Leisen et al., 2022) on *Arabidopsis thaliana* (Arabidopsis hereafter). In contrast to our previous findings on tomato, bean, maize and apple fruit, infection with the *hip1* strain led to significantly reduced lesion development, demonstrating that Hip1 contributes to *Botrytis* virulence on Arabidopsis specifically (Fig. 3A).

**Fig. 3:**
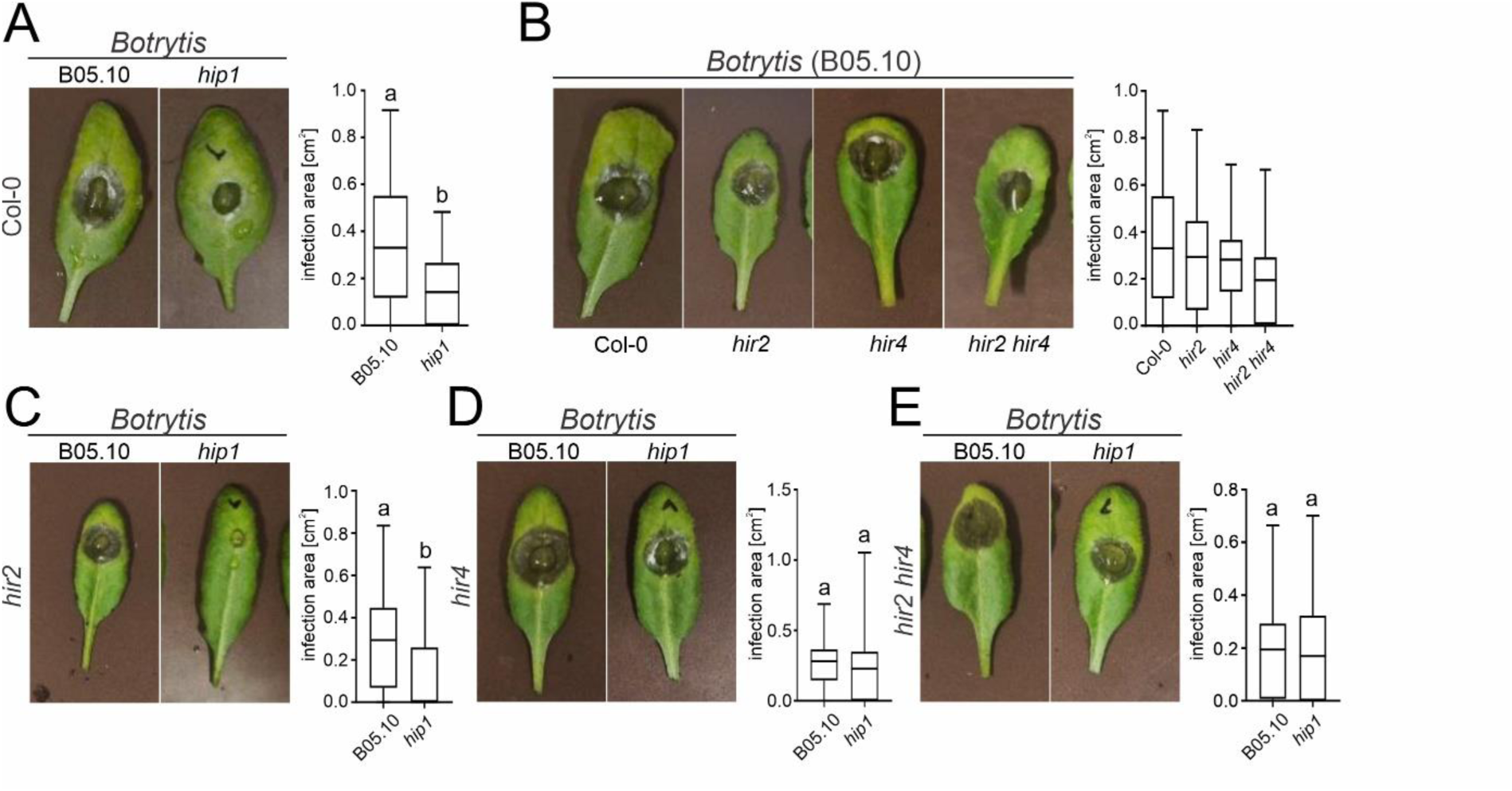
Infection tests using *Botrytis* wildtype and *hip1* knockout on different Arabidopsis genotypes. *Botrytis* wildtype (B05.10) and the *hip1* knockout mutant were used to infect Arabidopsis wildtype (Col-0), *hir2*, *hir4*, and the *hir2/hir4* double mutant. Representative pictures of 6-week-old Arabidopsis plants of the indicated genotype 3 days after infection with a 5 µl inoculum from a 1*10^6^ spores/ml suspension. A) *Botrytis* wildtype and *hip1* infection of Col-0; n=50, n=55. B) *Botrytis* infection of Col-0, *hir2*, *hir4* and *hir2 hir4*; n=50, n=58, n=47, n=51 C) *Botrytis* wildtype and *hip1* infection of hir2; n=58, n=52. D) *Botrytis* wildtype and *hip1* infection of *hir4*; n=47, n=47. E) *Botrytis* wildtype and *hip1* infection of *hir2 hir4*; n=51, n=52. Box limits in the graphs represent 25th-75th percentile, the horizontal line the median and whiskers minimum to maximum values. Different letters denote significant differences (one-way ANOVA with Tukey’s HSD post hoc test, *P* < 0.05). One-way ANOVA with Dunnett post hoc test was performed using Col-0 as control for (B), *\**p=0.05, ns: not significant. Scale bar is 0.5 cm. The experiment was performed 3 times with similar results.

To assess whether this virulence function depends on the proposed host targets of Hip1, we further analyzed disease development on *hir2*, *hir4*, and *hir2 hir4* mutant plants. In contrast to the findings reported for PEIE1 during *Sclerotinia* infection (Liu et al., 2024), loss of HIR proteins did not significantly affect susceptibility to infection *Botrytis* wild type (Fig. 3B). However, the virulence contribution of Hip1 was largely abolished in the hir4 mutant, whereas loss of HIR2 had no detectable effect (Fig. 3C–E). These findings suggest that Hip1-mediated virulence is genetically linked to HIR4.

### PEIE1 is a previously unrecognized phytotoxic protein

Hip1 has been previously described as highly phytotoxic protein that induces plant cell death (Jeblick et al., 2023). As PEIE1 and Hip1 are highly similar, we were interested whether PEIE1 also shares the phytotoxic property described for Hip1. Therefore, we cloned PEIE1 with signal peptide (sp) and without (nospPEIE1) and expressed them transiently in *N. benthamiana* by agroinfiltration.

Three days after transient expression, PEIE1 induced strong necrosis similar to that caused by Hip1, and this activity depended on the presence of a signal peptide (SP) (Fig. 4A and 4B; Supplementary Fig. 1). To further confirm the phytotoxic activity of PEIE1, recombinant PEIE1 was expressed in *E. coli* and affinity purified (4C). Infiltration of the purified protein into *N. benthamiana* leaves induced dose-dependent necrosis (4D), with visible symptoms observed at concentrations as low as 0.5 µM (Fig. 4E). To determine whether the phytotoxic activity of Hip1 and PEIE1 is conserved in Arabidopsis, purified recombinant proteins were infiltrated into Col-0 leaves. Consistent with our previous study, Hip1 induced little or no necrosis at concentrations up to 15 µM (Jeblick et al., 2023), and only weak necrosis became apparent at 20 µM (Supplementary Fig. 2B and C). Likewise, PEIE1 was only weakly phytotoxic in Arabidopsis, with detectable necrosis occurring only at the highest protein concentrations tested (Supplementary Fig. 2A). To assess whether HIR proteins contribute to this weak response, 20 µM Hip1 was infiltrated into the *hir2 hir4* double mutant and a HIR4 overexpression line. However, neither reduced nor elevated HIR abundance significantly altered Hip1-induced necrosis (Supplementary Fig. 2B and C). These findings demonstrate that, in contrast to *N. benthamiana*, both Alt a1-like proteins exhibit only limited phytotoxic activity in Arabidopsis and that this residual activity is independent of HIR abundance.

**Fig. 4:**
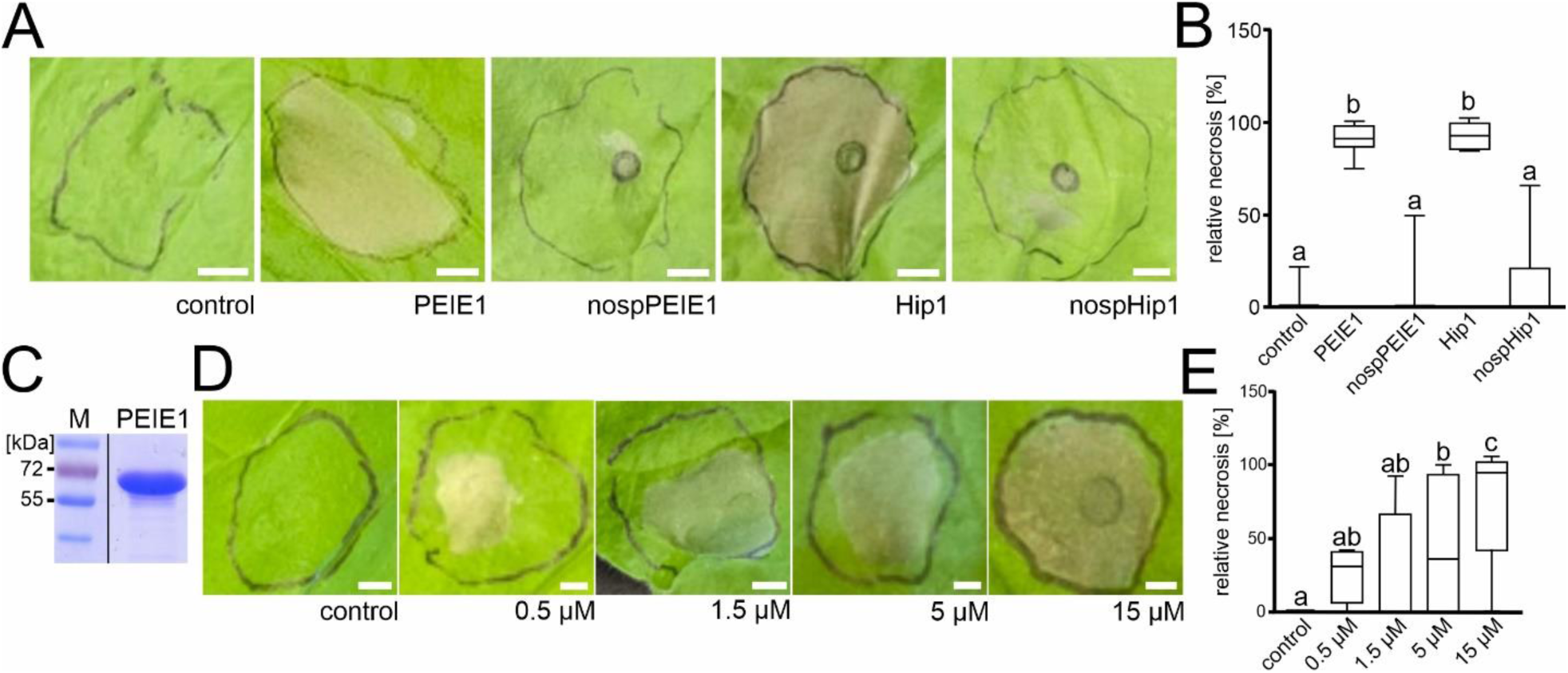
Extracellular Hip1 and PEIE1 cause cell death in tobacco leaves. A) PEIE1 and Hip1 were transiently expressed *in N. benthamiana* with and without signal peptide (nosp). Representative pictures are shown. Infiltration buffer was used as control. B) Necrotic area was quantified as the percentage of the infiltrated leaf area exhibiting visible necrosis. C) In *E. coli* expressed and purified PEIE1 protein. D) Infiltration of increasing PEIE1 concentrations in *N. benthamiana*. Dialysis buffer was used as control. Representative pictures are shown. E) Necrotic area was quantified as the percentage of the infiltrated leaf area exhibiting visible necrosis. Box limits in the graph represent 25th-75th percentile, the horizontal line the median and whiskers minimum to maximum values (n = 16-21 for (B) and n = 4-17 for (E)). Different letters denote significant differences (one-way ANOVA with Tukey’s HSD post hoc test, *P* < 0.05). Scale bars: 0.5 cm.

### Hip1-induced cell death requires the central immune regulator EDS1

To investigate the mechanism underlying the observed toxicity, the activity of Hip1 was examined in different *N. benthamiana* immune signaling mutants. As Hip1-induced cell death has previously been shown to occur independently of the cell surface co-receptors SOBIR1 and BAK1 (Jeblick et al., 2023), we focused on components of intracellular immune signaling, which are typically associated with effector-triggered immunity (ETI). In solanaceous plants, many coiled-coil nucleotide-binding leucine-rich repeat receptors (CNLs) signal through the NLR REQUIRED FOR CELL DEATH (NRC) helper NLR network (Adachi et al., 2019; Wu et al., 2017). However, Hip1-induced cell death was unaffected in two independent *nrc2 nrc3 nrc4* triple mutant lines, indicating that NRC-dependent CNL signaling is not required for Hip1 activity (Supplementary Fig. 3). We therefore investigated the contribution of the EDS1-dependent immune signaling pathway. EDS1 is a central regulator of intracellular immunity, forming heterodimers with either PAD4 or SAG101 to transduce signals from activated TIR-NLR (TNL) receptors, thereby promoting defence responses and host cell death (Dongus and Parker, 2021) (Fig. 5A). Using purified recombinant protein (Jeblick et al., 2023), we assessed Hip1-induced cell death in a series of published *eds1*, *pad4*, and *sag101a/sag101b* single and higher-order *N. benthamiana* mutant lines (Ordon et al., 2021; Ordon et al., 2017; Prautsch et al., 2023; Lapin et al., 2019; Gantner et al., 2019). Notably, all genotypes lacking EDS1 exhibited a marked reduction in necrosis development compared with the wild type (Figure 5B). PEIE1-induced cell death likewise depended on EDS1 but not on SOBIR1/SOBIR-like receptors (Supplementary Fig. 4), indicating that both Alt a1-like proteins engage a common immune signaling pathway involving EDS1.

**Fig. 5:**
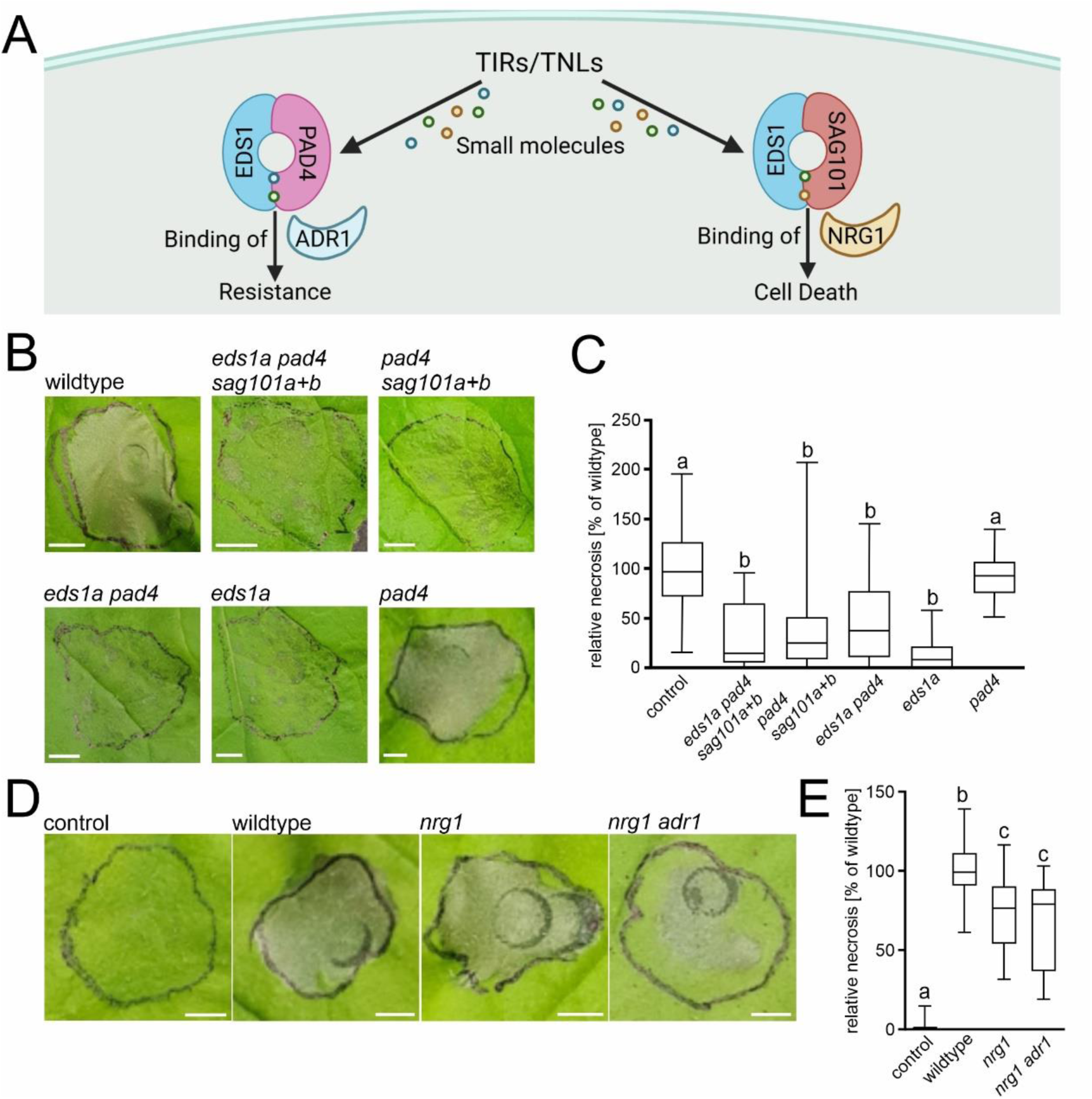
Hip1-induced cell death requires EDS1-dependent immune signaling. A) Schematic overview of TIR/TNL-mediated immune signaling, illustrating the EDS1–PAD4–ADR1 branch associated primarily with pathogen resistance and the EDS1–SAG101–NRG1 branch associated with hypersensitive cell death (created with BioRender). B) Purified recombinant Hip1 protein produced in *E. coli* was infiltrated into wild type and immune-deficient *N. benthamiana* mutant lines lacking components of the EDS1 signaling pathway. C) Necrotic area was quantified as the percentage of the infiltrated leaf area showing visible necrosis and in relation to the wt symptoms. D) Phenotypic symptoms and E) quantification of relative necrosis following infiltration of Hip1 protein into wildtype, *nrg1*, and *nrg1 adr1* double-knockout mutant leaves. Box limits represent the 25th–75th percentiles, the horizontal line indicates the median, and whiskers represent the minimum and maximum values (n = 26–33 for (B) and n = 16-31 for (C)). Different letters denote significant differences (one-way ANOVA with Tukey’s HSD post hoc test, *P* < 0.05). Scale bars: 0.5 cm.

### Hip1 engages EDS1-dependent immunity through downstream signaling

To identify immune signaling components acting downstream of EDS1, purified recombinant Hip1 protein was infiltrated into *N. benthamiana nrg1* and *nrg1 adr1* mutant plants (Figure 5C). In Arabidopsis, EDS1–PAD4 and EDS1–SAG101 complexes preferentially engage distinct downstream signaling branches through the helper NLR families ADR1 and NRG1, respectively, which contribute differentially to TNL-mediated immunity and cell death (Lapin et al., 2019; Huang et al., 2022). Similar helper NLR networks have been described in *N. benthamiana*, which contains an expanded family of ADR1 homologues (Ordon et al., 2021). Hip1-induced necrosis was significantly reduced in both the *nrg1*-single mutant (Ordon et al., 2021) and the *nrg1 adr1* double mutant (Prautsch et al., 2023) compared with wild-type plants (Figure 5C), demonstrating that helper NLR signaling contributes to Hip1-induced cell death.

## Discussion

In *Botrytis*, diverse mechanisms of host cell death induction have been described, including the pore-forming activity of necrosis– and ethylene-inducing proteins (NLPs), enzymatic degradation of the plant cell wall by endopolygalacturonases and other glycoside hydrolases, the necrosis-inducing activity of cerato-platanins, and the intracellular activity of the recently identified effector Crh1 (Have et al., 1998; Kars et al., 2005; Frías et al., 2011; Bi et al., 2021; Seidl and van den Ackerveken, 2019; Leisen et al., 2022; Liang et al., 2026). Although these proteins employ distinct biochemical mechanisms and are perceived through different host pathways, they ultimately converge on the induction of host cell death, a process that facilitates colonization by necrotrophic pathogens. Compared with many other CDIPs, however, the signaling mechanism underlying the activity of Hip1 has remained largely unresolved). Hip1 has previously been characterized as a potent phytotoxic protein acting as a PAMP-like elicitor (Jeblick et al., 2023). Here, we identify Hip1 as a structural and functional homolog of the recently described *Sclerotinia* effector PEIE1 and demonstrate an unexpected requirement for the central immune regulator EDS1 and its downstream helper NLR network for Hip1-induced cell death.

Hip1 and PEIE1 are closely related members of the fungal Alt a1-like protein family. Both proteins share the characteristic β-barrel fold originally described for the major allergen Alt a1 from *Alternaria alternata*, a structural scaffold conserved among proteins from numerous plant-pathogenic fungi(Garrido-Arandia et al., 2016; Jeblick et al., 2023; Chruszcz et al., 2012). Consistent with their high sequence and predicted structural similarity, Hip1 and PEIE1 both interacted with the plasma membrane-localized HIR protein HIR4. In contrast, the related *Botrytis* protein Hip2, despite adopting the same predicted Alt a1-like fold, failed to interact with HIR4, suggesting that HIR recognition depends on conserved surface-exposed residues rather than the common structural scaffold itself. These observations are consistent with the recent work of Liu et al. (2024), who demonstrated that PEIE1 associates with HIR proteins and interferes with HIR oligomerization during *Sclerotinia* infection.

Despite their interaction with Arabidopsis HIR proteins, both Hip1 and PEIE1 exhibited only weak phytotoxic activity when infiltrated into Arabidopsis leaves, whereas both proteins efficiently induced necrosis in *N. benthamiana*. Moreover, altering HIR abundance through loss of *HIR2* and *HIR4* or overexpression of *HIR4* did not significantly affect Hip1-induced necrosis. These observations indicate that cell death induction is independent of HIR binding. and suggest that additional host-specific immune components determine whether recognition of Alt a1-like proteins culminates in a necrotic response. Such host-dependent differences are well documented for microbial elicitors and reflect interspecific variation in immune receptor repertoires, receptor complex formation, and downstream signaling capacity (Boutrot and Zipfel, 2017; Ngou et al., 2022).

PEIE1 itself exhibits strong phytotoxic activity in *N. benthamiana*, comparable to Hip1. While Liu et al. (2024) identified PEIE1 as an immunosuppressive effector that dampens HIR-mediated immunity, its capacity to induce host cell death was not investigated. Our results therefore reveal an additional functional dimension of PEIE1 and suggest that its biological activity depends strongly on host context. The robust necrosis observed in *N. benthamiana*, together with the lack of obvious toxicity reported in *Arabidopsis*, is consistent with the species-specific perception observed for numerous fungal PAMP-like proteins and CDIPs (Frías et al., 2011; Bi et al., 2021). Thus, rather than acting solely as an immune suppressor, PEIE1 appears capable of activating immune-mediated cell death under appropriate conditions.

In contrast to PEIE1, whose virulence function was associated with altered susceptibility of *hir* mutants during *Sclerotinia* infection (Liu et al., 2024), loss of *HIR2* or *HIR4* did not significantly alter susceptibility to wild-type *Botrytis cinerea*. Nevertheless, the contribution of *Hip1* to fungal virulence was specifically dependent on plant *HIR4*, as the *Botrytis hip1* deletion strain no longer showed reduced virulence in the Arabidopsis *hir4* background. This genetic interaction supports our biochemical data identifying HIR4 as a Hip1-interacting protein and suggests that HIR4 contributes specifically to Hip1 function during infection rather than acting as a general determinant of *Botrytis* susceptibility. The absence of a comparable effect in *hir2* mutants further indicates that, although Hip1 can associate with both HIR2 and HIR4, these closely related proteins are not functionally equivalent during fungal infection.

Recent studies have considerably expanded our understanding of HIR protein function and provide an intriguing framework for interpreting these observations. HIR proteins belong to the plant-specific SPFH (stomatin/prohibitin/flotillin/HflK/C) protein family and accumulate within specialized plasma membrane nanodomains during immune activation (Qi et al., 2011; Hdedeh et al., 2025). While initially identified as proteins associated with the hypersensitive response (Jung and Hwang, 2007), emerging evidence indicates that HIR2 functions as a central organizer of immune signaling complexes, dynamically recruiting receptors and signaling proteins during both pattern-triggered and intracellular immunity (Weber et al., 2026). Rather than acting as passive structural components, HIR proteins appear to organize membrane nanodomains that integrate extracellular perception with downstream immune signaling (Jaillais and Ott, 2020). The interaction of both Hip1 and PEIE1 with HIR proteins therefore suggests that these fungal proteins converge on a central immune signaling hub rather than targeting a single receptor.

The most unexpected finding of this study is the requirement of EDS1 for Hip1-induced cell death. EDS1 is widely recognized as the central signaling hub of TIR-NLR (TNL)-mediated immunity, where it forms heterodimers with either PAD4 or SAG101 to activate the helper NLR families ADR1 and NRG1, respectively (Lapin et al., 2019; Dongus and Parker, 2021). Consistent with this model, Hip1-induced cell death was strongly reduced not only in *eds1* mutants but also in *adr1*, *nrg1*, and *adr1 nrg1* mutant plants, indicating that Hip1-induced necrosis depends on the canonical EDS1 helper NLR signaling network. Although the EDS1–PAD4–ADR1 and EDS1–SAG101–NRG1 branches exhibit partial functional specialization, increasing evidence suggests substantial crosstalk and genetic redundancy between these pathways (Wu et al., 2019; Gao et al., 2019; Sun et al., 2021). Our findings therefore extend the known role of this signaling network by demonstrating that it can be engaged by an extracellular fungal CDIP.

Together, our findings support a model in which Hip1 and PEIE1 possess at least two separable biological activities. First, both proteins interact with the plasma membrane-associated HIR proteins, which likely functions as organizers of immune signaling nanodomains (Weber et al., 2026). In *Sclerotinia*, PEIE1 has been shown to interfere with HIR oligomerization and suppress host immunity (Liu et al., 2024), suggesting that HIR binding may represent a conserved molecular function of this fungal protein family. Second, both proteins are capable of activating immune-mediated cell death, although the outcome is strongly host dependent. Whereas Hip1 and PEIE1 trigger robust EDS1-dependent necrosis in *N. benthamiana*, both proteins exhibit only weak phytotoxicity in Arabidopsis, despite interacting with Arabidopsis HIR proteins. Although these activities initially appear contradictory, they are not necessarily mutually exclusive and may occur at different stages of infection or represent host-dependent outcomes of HIR perturbation. We therefore propose that HIR association represents an upstream event that can lead to distinct biological outputs depending on the host species and immune signaling context. During natural infection, Hip1 may contribute to fungal virulence through modulation of HIR-associated immune signaling, while strong necrotic responses become apparent only in hosts that possess the appropriate downstream signaling machinery.

In conclusion, our study identifies Hip1 as a structural and functional homolog of the *Sclerotinia* effector PEIE1 and reveals an unexpected link between Alt a1-like fungal proteins and the plant immune regulators HIR and EDS1. Although Hip1 contributes to *Botrytis* virulence in Arabidopsis, both Hip1 and PEIE1 exhibit only weak phytotoxic activity in this host while inducing robust cell death in *N. benthamiana*. Together with the requirement for EDS1 and helper NLR signaling, these findings suggest that these Alt a1-like fungal proteins engage conserved immune signaling components but that the resulting biological outcome is strongly shaped by the host-specific immune signaling network. Our work therefore provides a framework for understanding how structurally conserved fungal proteins can exert distinct effects on immunity and disease development in different plant species.

## Materials & methods

### Plant Material and Growth Conditions

*N. benthamiana* was grown for 4–6 weeks on soil at 23°C under long-day conditions (14-h light/10-h dark) for transient expression and leaf infiltration. *eds1* (Ordon et al., 2017), PAD4 and SAG101 mutants (Gantner et al., 2019), *eds1 pad4 sag101a sag101b* (Lapin et al., 2019), *nrg1* (Ordon et al., 2021) and *nrg1 adr1* (Prautsch et al., 2023) mutants as well as *nrc2 nrc3 nrc4* #431 and *nrc2 nrc3 nrc4* #551 (Wu et al., 2020) has been published before. For infection and infiltration tests, Arabidopsis (*A. thaliana*) ecotype Col-0 was grown as wild type under short-day regime (8-h light/16-h dark) for 5-6 weeks at 22°C. The following mutants were published previously (Liu et al., 2024): *hir2*, *hir4*, *hir2 hir4*, oxAtHIR4-8 (HIR4^OE^).

### Cultivation of *Botrytis* and infection tests

*Botrytis cinerea* wild type (B05.10) was used for infection tests. The hip1 gene knockout strain has been published before (Leisen et al., 2022). Cultivation of *Botrytis* was performed as described previously (Müller et al., 2018). Infection tests were carried out using and 1 × 10^6^ spores mL^−1^, pregerminated for 6 h in semi-solid medium (0.35% agar). Resulting lesions were documented after 72 h and lesion size (area) was quantified using the freehand tool in ImageJ.

### Recombinant plasmid construction

Coding sequence of PEIE1 was synthesized. Full-length PEIE1 was generated by a Phusion PCR protocol using the corresponding primer pairs. Destination vector and PCR fragments for everything except mbSUS experiments were digested with the Type IIS restriction enzyme Bsa1 at the indicated sites (Supplementary Table) and ligation reactions were performed using T4 DNA ligase (New England BioLabs). For transient expression, a GreenGate cloning system was employed (Lampropoulos et al., 2013), pGGZ003 was used as destination vector, pGGZ003 harbors a cauliflower mosaic virus (CMV) 35S promoter, an N-terminal signal peptide (SP), an RCBS terminator, an HA-tag for immunological detection, and a spectinomycin resistance cassette. For recombinant protein expression, the Hip1 coding sequence lacking the signal peptide was amplified and cloned into the pET28a (+) expression vector, containing an N-terminal His-tag fusion as well. The PEIE1 coding sequence has been cloned into the pCold expression vector, containing a trigger factor tag for enhanced solubility. All gene constructs were verified by Sanger sequencing (Seq-It, Germany). All primers used are shown in the table below. Gene constructs were verified by sequencing (Eurofins Genomics, Light Run, Germany).

For interaction experiments, coding sequences were cloned into suitable entry and destination vector using Gateway cloning.

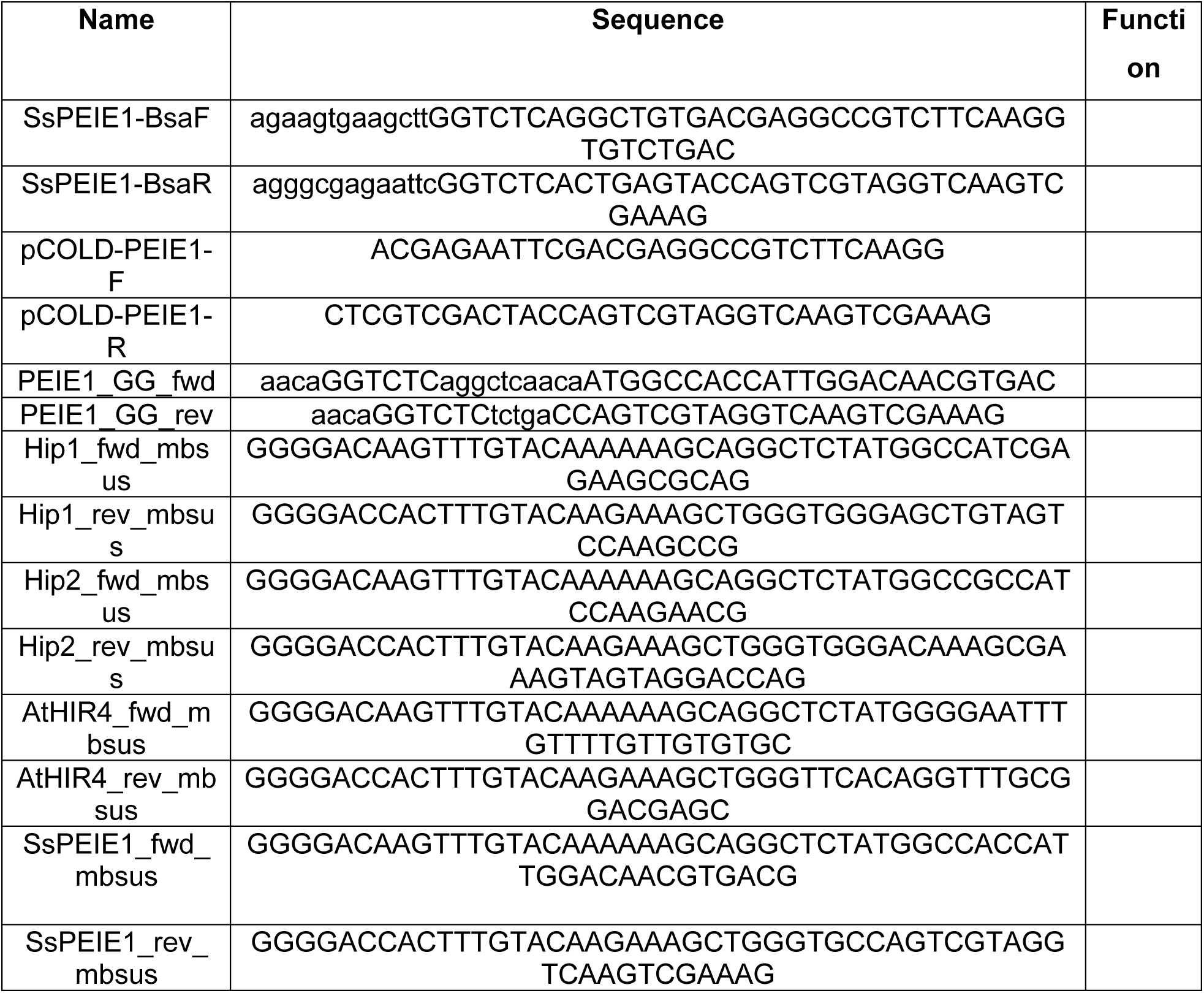

### Sequence alignments and structural modeling

Sequence alignments were carried out using clustalW. To structurally compare Hip1 and PEIE1 3D protein modelling was performed using AlphaFold. The obtained predicted structural model allowed the visualization of the spatial organization of the protein as well as the model confidence using the predicted Local Distance Difference scores.

### Transient expression in *N. benthamiana*

For transient expression in *N. benthamiana*, the *A. tumefaciens* (strain GV3101::mp90) was used as described previously (Müller et al., 2026), without using the p19 helper strain. After transient expression (3 days) proteins were extracted from *N. benthamiana* leaf disks (20 mm diameter) as described before (Müller et al., 2026).

### Recombinant protein expression

*Escherichia coli* strain T7 shuffle (New England Biolabs, USA) was used to express Hip1 and PEIE1. Induction of protein expression was enabled by 0.5 mM isopropyl β-d-1-thiogalactopyranoside (IPTG) and conducted at 20°C overnight. Solubility was checked using B-PER™ Reagent (Thermo scientific, USA). After bacterial lysis, protein purification was performed via immobilized metal affinity chromatography using His/Ni beads (Roth, Germany) as resin. Elution buffer has been exchanged with a 3x 1h dialysis in 50 mM phosphate buffer with 150 mM NaCl (pH 6).

### Statistical analysis

Analysis was carried out using the GraphPad Prism 9 software. The detailed statistical method employed is provided in the respective figure legends. All experiments were carried out at least three times. Box limits in the graphs represent 25th to 75th percentile, the horizontal line the median and whiskers minimum to maximum values.

## Acknowledgements

We would like to thank Johannes Stuttmann, Jane Parker and Sophien Kamoun for providing published plant material, Sabrina Kaiser for technical help with the mbSUS experiments and Birgit Kemmerling for critical reading of the manuscript. This work was supported by grants from the *BioComp* research initiative (Rhineland-Palatinate, Germany) and the German research foundation (DFG; SCHE 1836/5-1) to MH and DS.

## Contributions

TM, MM, CC, SK, BK and DS performed and analysed experiments. TM and DS designed the figures and performed statistical analysis. MH and DS conceived the study and DS wrote the manuscript. All authors saw and commented on the manuscript.

## Corresponding author

Correspondence to David Scheuring.

