## Supplementary Figures for "Two homologous Alt a1-like fungal proteins possess dual activities in HIR-associated immune signaling and EDS1-dependent cell death"

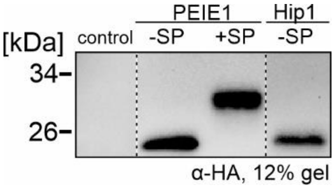


**Supplementary Fig. 1: Confirmation of transient expression via immunoblotting.** An HA-antibody directly coupled to HRP was used for detection.

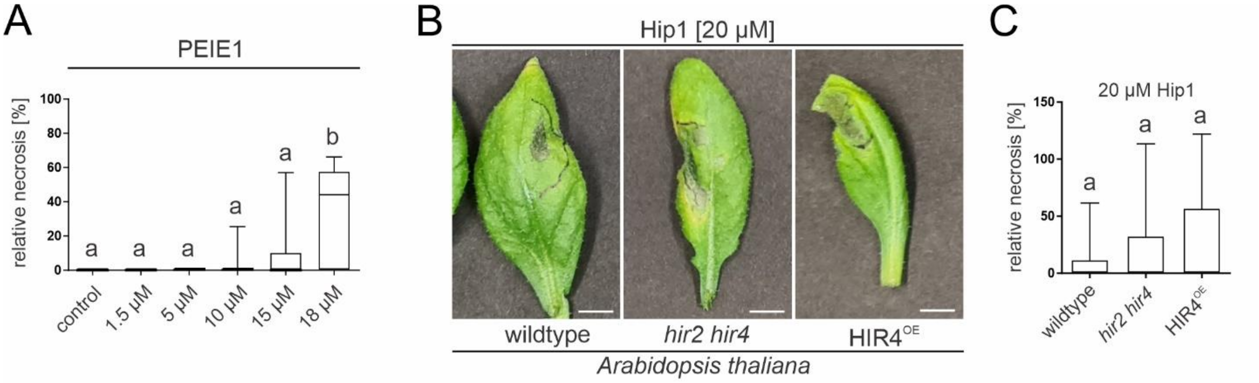


**Supplementary Fig. 2. PEIE1 and Hip1 exhibit only weak phytotoxicity at high protein concentrations in *Arabidopsis*, independent of HIR.** A) *Arabidopsis thaliana* wild-type (Col-0) leaves were infiltrated with increasing concentrations of PEIE1 protein. Cell death was quantified 48 h after infiltration. B) Leaves of *A. thaliana* wildtype, the *hir2 hir4* double mutant, and an HIR overexpression line were infiltrated with 20 µM Hip1 protein, and cell death was quantified 48 h after infiltration. C) The necrotic area was quantified as the percentage of the infiltrated leaf area displaying visible necrosis. Box plots show the 25th and 75th percentiles, with the median indicated by the horizontal line; whiskers represent the minimum and maximum values. Different letters denote significant differences (one-way ANOVA followed by Tukey's HSD post hoc test, $P < 0.05$). Scale bars, 0.5 cm. Different letters denote significant differences (one-way ANOVA with Tukey's HSD post hoc test, $P < 0.05$). Scale bars: 0.5 cm.

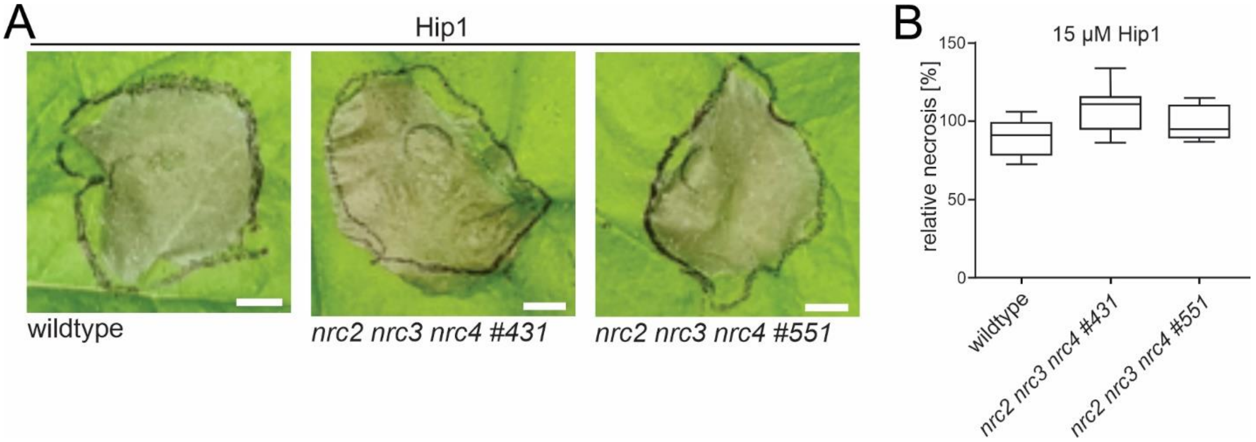


**Supplementary Fig. 3 Hip1-induced cell death is maintained in *nrc2 nrc3 nrc4* triple mutants.** *N. benthamiana* wild-type and two independent *nrc2 nrc3 nrc4* mutant lines (#431 and #551) infiltrated with 15 µM Hip1 protein. Cell death was quantified 48h after infiltration. A) Representative pictures are shown. B) Necrotic area was quantified as the percentage of the infiltrated leaf area exhibiting visible necrosis. Box limits in the graph represent 25th-75th percentile, the horizontal line the median and whiskers minimum to maximum values. Scale bars: 0.5 cm.

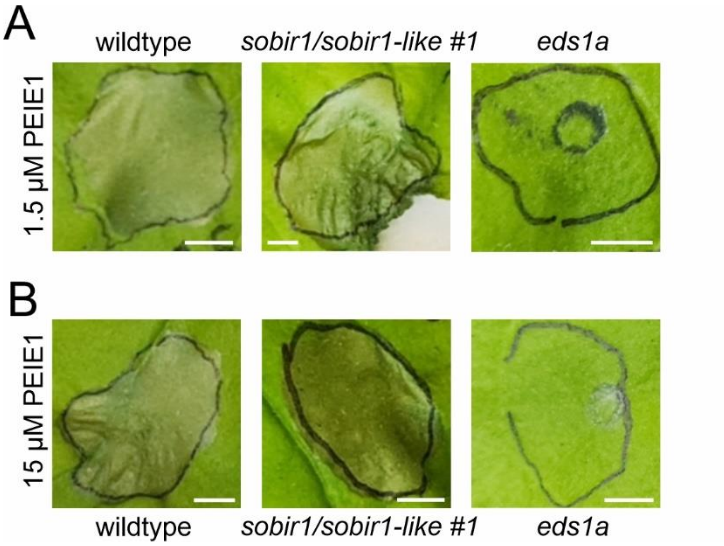


**Supplementary Fig. 4: PEIE1-induced cell death is independent of SOBIR1/SOBIR1-lie but depends on EDS1.** *N. benthamiana* wild-type and *sobir1/sobir1-like* as well as *eds1a* were infiltrated with PEIE1 protein and cell death quantified 48h after infiltration. A) 1.5 µM PEIE1 was infiltrated. B) 15 µM PEIE1 was infiltrated.
